# Data from Entomological Collections of Aedes (Diptera: Culicidae) in a post-epidemic area of Chikungunya, City of Kinshasa, Democratic Republic of Congo

**DOI:** 10.1101/2023.09.20.558445

**Authors:** Victoire Nsabatien, Josue Zanga, Fiacre Agossa, Nono Mvuama, Maxwell Bamba, Osée Mansiangi, Leon Mbashi, Vanessa Mvudi, Glodie Diza, Dorcas Kantin, Narcisse Basosila, Hyacinthe Lukoki, Arsene Bokulu, Christelle Bosulu, Erick Bukaka, Jonas Nagahuedi, Jean Claude Palata, Emery Metelo

## Abstract

Arbovirus epidemics (*e*.*g*. Chikungunya, dengue, West Nile, Yellow Fever, and Zika), are a growing threat in Africa in areas where *Aedes* (*Ae*.) *aegypti* and *A. albopictus* are present.

The lack of complete sampling of these two vectors limits our ability to understand their propagation dynamics in areas at risk from arboviruses. Here, we describe for the first time the geographical distribution of two arbovirus vectors (*Ae. aegypti* and *Ae. albopictus*) in a chikungunya post-epidemic zone in the provincial city of Kinshasa, Democratic Republic of Congo between 2020 and 2022. In total 6,943 observations were reported using larval capture and human capture on landing methods. These data are published in the public domain as a Darwin Core archive in the Global Biodiversity Information Facility. The results of this study potentially provide important information for further basic and advanced studies on the ecology and phenology of these vectors, as well as on vector dynamics after an epidemic period.

**Subject Areas:** Ecology, Biodiversity, Taxonomy

**Data description:** 

## Background and context

Arbovirus epidemics in general, and Chikungunya in particular, are a growing threat in Africa in areas where *Aedes* (Ae.) *aegypti* and *albopictus* are present (1,2,3,4).

The lack of complete sampling of these two vectors limits our ability to understand their propagation dynamics in areas at risk from arboviruses. Thus, studies such as ours have contributed to understanding the dynamics of these vectors.

Here, we describe for the first time the geographical distribution of two arbovirus vectors (*Ae. aegypti* and *Ae. albopictus*) in a chikungunya post-epidemic zone in the provincial city of Kinshasa, Democratic Republic of Congo between 2020 and 2022.

## Methods

### General spatial coverage

The study was carried out in the Vallée de la Funa (Figure 1), a forest gallery located in the hilly area of the commune of Mont Ngafula. It is located within a secondary forest island dominated by *Millettia laurentii* and *Pentaclethra eetveldeana*.

**Fig 1.**
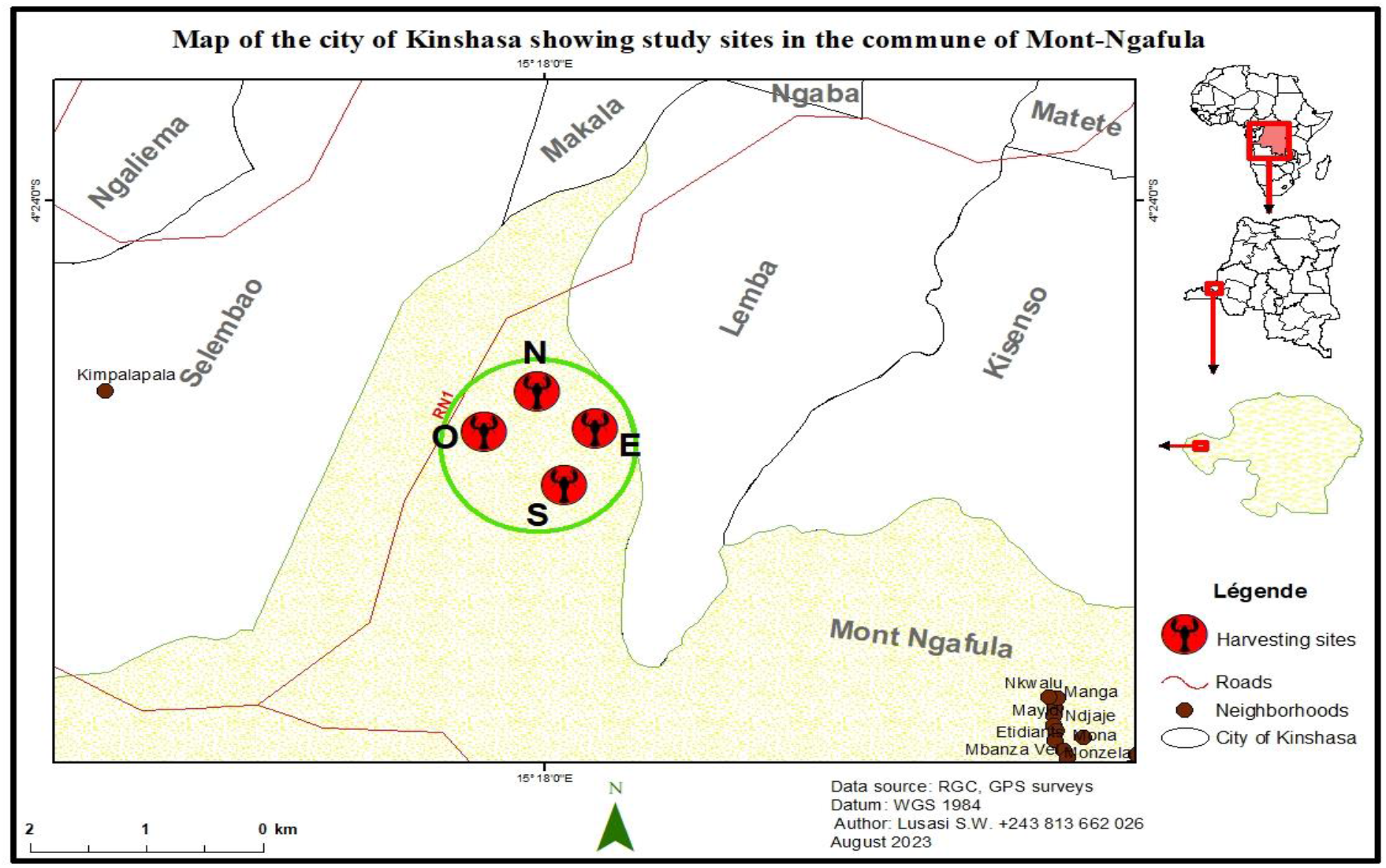
Map of the Funa Valley, showing entomological collection points (North, South, East and West).

The forest gallery is located between latitude 4°25’06.17’’S and longitude 15°18’05.77’’E, with a surface area of around 45 hectares. Average annual rainfall is around 1,095 mm. However, rainfall in Kinshasa is abundant and unevenly distributed throughout the year. The highest volume of precipitation occurs in November, with an average of 268.1 mm, and the lowest volume occurs in July, with an average of 0.7 mm. And the relative annual average humidity is 79%.

### Coordinates

4°26’22.27’’S and 4°16’40.59’’S Latitude; 15°14’16.93’’E and 15°26’38.51’’E Longitude

### Mosquito collection

The general taxonomic coverage description for this work is the Culicidae Family, specifically *Aedes aegypti* (commonly known as Yellow fever mosquito, NCBI:txid7159) and *Aedes albopictus* (Commonly known as the Asian tiger mosquito or moustique tigre in French, NCBI:txid7160)

Two sampling techniques were used to collect immature and adult stages of *Aedes* spp. between : January 5, 2020 - December 20, 2022. Whatever the collection method, morphological identification of adult mosquitoes was carried out using the taxonomic keys (5).

### Larval collection

Immature stages of *Aedes* spp. were sampled from various breeding sites (abandoned jars, tires, cans and other containers) using the dipping technique. They were filtered and stored in labelled jars, then transported to the insectarium of the Laboratory of Bioecology and Vector Control (BIOLAV) for rearing to adulthood.

### Human capture on landing 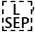

This method was used to collect *Aedes* spp. adults during the day. At the study site, mosquito collections were carried out both indoors and outdoors, with a first group of collectors working from 7:00 am to 12:00 pm and replaced by a second group from 12:00 pm to 7:00 pm. At each collection point, a bare-legged, barefoot volunteer served as bait, collecting mosquitoes using hemolysis tubes. Specimens of *Aedes* spp. were collected, released in cages and transported to the insectarium.

### Quality control description

Mosquitoes were identified using keys available in the literature (5) by an entomologist experienced in the identification of Central African *Aedes*.

Once digitized the data has also been validated using the validator available in Global Biodiversity Information Facility (GBIF).

## Results

A total of 6,943 observations (summarized in TABLE 1) were reported using larval capture and human capture on landing methods during 2020 and 2022. The human landing catch collecting method was responsible for 4,638 adult mosquitoes collected, while the larval sampling yielded 2,305 individuals. *Aedes albopictus* is the most abundant species in the studied region with 6,358 individuals and *A*.*aegypti* with 578. A total of 7 individuals could not be identified to species level.

**Table 1.**
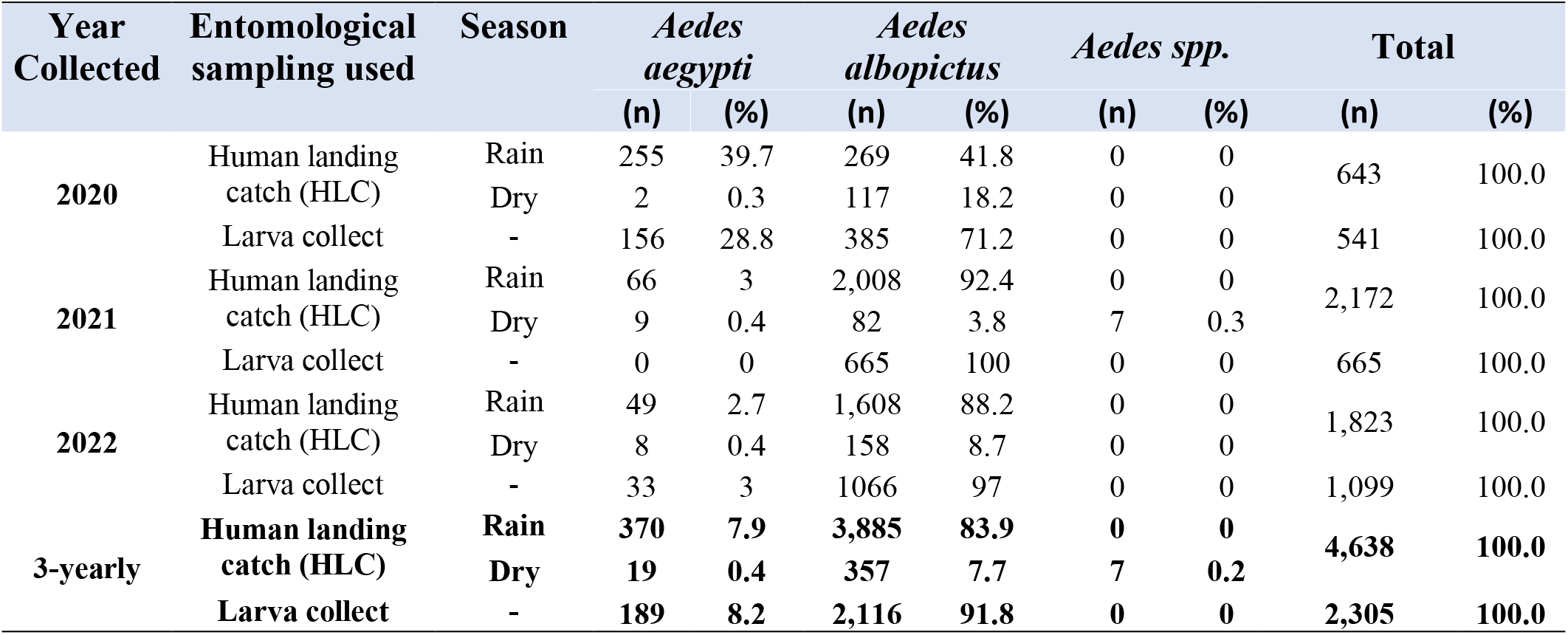
Summary of results obtained using two entomological sampling techniques after three years of study in a Chikungunya post-epidemic area.

| Year Collected | Entomological sampling used | Season | *Aedes aegypti* |  | *Aedes albopictus* |  | *Aedes spp.* |  | Total |  |
| --- | --- | --- | --- | --- | --- | --- | --- | --- | --- | --- |
|  |  |  | (n) | (%) | (n) | (%) | (n) | (%) | (n) | (%) |
| 2020 | Human landing catch (HLC) | Rain | 255 | 39.7 | 269 | 41.8 | 0 | 0 | 643 | 100.0 |
|  |  | Dry | 2 | 0.3 | 117 | 18.2 | 0 | 0 |  |  |
|  | Larva collect | - | 156 | 28.8 | 385 | 71.2 | 0 | 0 | 541 | 100.0 |
| 2021 | Human landing catch (HLC) | Rain | 66 | 3 | 2,008 | 92.4 | 0 | 0 | 2,172 | 100.0 |
|  |  | Dry | 9 | 0.4 | 82 | 3.8 | 7 | 0.3 |  |  |
|  | Larva collect | - | 0 | 0 | 665 | 100 | 0 | 0 | 665 | 100.0 |
| 2022 | Human landing catch (HLC) | Rain | 49 | 2.7 | 1,608 | 88.2 | 0 | 0 | 1,823 | 100.0 |
|  |  | Dry | 8 | 0.4 | 158 | 8.7 | 0 | 0 |  |  |
|  | Larva collect | - | 33 | 3 | 1066 | 97 | 0 | 0 | 1,099 | 100.0 |
| 3-yearly | **Human landing catch (HLC)** | **Rain** | **370** | **7.9** | **3,885** | **83.9** | **0** | **0** | **4,638** | **100.0** |
|  |  | **Dry** | **19** | **0.4** | **357** | **7.7** | **7** | **0.2** |  |  |
|  | **Larva collect** | **-** | **189** | **8.2** | **2,116** | **91.8** | **0** | **0** | **2,305** | **100.0** |

### Re-use potential

The results of this study will provide important information for further basic and advanced studies on the ecology and phenology of these vectors, as well as on vector dynamics after an epidemic period. Given that these vectors have been responsible for a considerable burden of disease during the various Chikungunya epidemic periods in DR. Congo, the results of these entomological surveys can help national policy-makers to target appropriate interventions and allocate resources to national vector control programs. But also to explain why existing interventions are not working as intended.

## Data Availability

The data supporting this article are published through the Integrated Publishing Toolkit (IPT) of University of Kinshasa and are available via GBIF under a CC0 waiver [1]. https://doi.org/10.15468/q53h26

## Dataset description

**Object name:** Darwin Core Archive Data from Entomological Collections of Aedes (Diptera: Culicidae) in a post-epidemic area of Chikungunya, City of Kinshasa, Democratic Republic of Congo

**Character encoding:** UTF-8

**Format name:** Darwin Core Archive format

**Format version:** 1.0

**Distribution:** https://cloud.gbif.org/africa/archive.do?r=aedes_drc

**Publication date of data:** 2023-06-30

**Language:** English

**Licences of use:** [Public Domain (CC0 1.0)](https://creativecommons.org/publicdomain/zero/1.0/legalcode)

**Metadata language:** English

**Date of metadata creation:** 2023-06-30

**Hierarchy level:** Dataset

## Acknowledgements

The authors would like to thank the Bioecology and Vector Control Laboratory at Kinshasa School of Public Health for providing the facilities to colonize the larval stages of Aedes until emergence, and the instruments required for mosquito identification. In addition, many thanks to Dear Willy LUSASI for his willingness to design the map of the Funa valley, showing the entomological collection points for this study. We would also like to thank the communities of the Funa Valley for allowing us into their homes to collect samples.

## References

1. Kamgang B., Nchoutpouen E., Simard F. & Paupy C (2012). Notes on the blood-feeding behavior of Aedes albopictus (Diptera: Culicidae) in Cameroon. Parasite. Vecteurs. 5:57.

2. Zahouli J., Koudou B., Müller P., Malone D., Tano Y. & Utzinger J (2017). Urbanization is a main driver for the larval ecology of Aedes mosquitoes in arbovirus-endemic settings in south-eastern Côte d’Ivoire. PLos Negl Trop. Dis. 11

3. Fritz M., Taty Taty R., Portella C., Guimbi C., Mankou M., Leroy EM. & Becquart P (2019). Re-emergence of chikungunya in the Republic of the Congo in 2019 associated with a possible vector-host switch. International Journal of Infectious Diseases 84; 99–101.

4. Proesmans S., Katshongo F., Milambu J., Fungula B., Mavoko H., Ahuka-Mundeke S. et al. (2019). Dengue and chikungunya among outpatients with acute undifferentiated fever in Kinshasa, Democratic Republic of Congo: a crosssectional study. PLoS Negl Trop Dis. 13(9):1–16.

5. Huang Y & Ward R (1982). A pictorial key for the identification of the mosquitoes associated with yellow fever in Africa. Mosq. Syst.(1981) 13: 138149.

